# The density of regulatory information is a major determinant of evolutionary constraint on non-coding DNA in *Drosophila*

**DOI:** 10.1101/2023.07.21.550043

**Authors:** Gonzalo Sabarís, Daniela M. Ortíz, Ian Laiker, Ignacio Mayansky, Sujay Naik, Giacomo Cavalli, David L. Stern, Ella Preger-Ben Noon, Nicolás Frankel

**Author notes:** equal contribution (Gonzalo Sabarís, Daniela M. Ortíz, Ian Laiker).

## Abstract

The density and distribution of regulatory information in non-coding DNA of eukaryotic genomes is largely unknown. Evolutionary analyses have estimated that ∼60% of nucleotides in intergenic regions of the *D. melanogaster* genome is functionally relevant. This estimate is difficult to reconcile with the commonly accepted idea that enhancers are compact regulatory elements that generally encompass less than 1 kilobase of DNA. Here, we approached this issue through a functional dissection of the regulatory region of the gene *shavenbaby* (*svb*). Most of the ∼90 kilobases of this large regulatory region is highly conserved in the genus *Drosophila*, though characterized enhancers occupy a small fraction of this region. By analyzing the regulation of *svb* in different contexts of *Drosophila* development, we found that the regulatory architecture that drives *svb* expression in the abdominal pupal epidermis is organized in a dramatically different way than the information that drives *svb* expression in the embryonic epidermis. While in the embryonic epidermis *svb* is activated by compact and dispersed enhancers, *svb* expression in the pupal epidermis is driven by large regions with enhancer activity, which occupy a great portion of the *svb cis*-regulatory DNA. We observed that other developmental genes also display a dense distribution of putative regulatory elements in their regulatory regions. Furthermore, we found that a large percentage of conserved non-coding DNA of the *Drosophila* genome is contained within putative regulatory DNA. These results suggest that part of the evolutionary constraint on non-coding DNA of *Drosophila* is explained by the density of regulatory information.

## Introduction

Genomic regions that encode the information for gene regulation have been studied intensely for decades (Davidson 2010; Schaffner 2015; Moore et al. 2020). A clear picture has emerged from these analyses, where gene expression is controlled through the combinatorial binding of transcription factors (TFs) to regulatory elements such as transcriptional enhancers (Zinzen et al. 2009). Enhancers contain arrangements of transcription factor binding sites (TFBSs) that promote transcription at defined spatio-temporal patterns (Levine 2010). Decades of work have shown that enhancers play an important role in evolutionary change (Carroll 2008; Hill et al. 2021), and that their malfunction can cause disease (Claringbould and Zaugg 2021).

It is generally assumed that the regulatory information for driving an expression pattern is mostly contained within DNA regions shorter than 1 kilobase. That is, the prevailing view of gene regulation envisions enhancers as compact regulatory elements (Levine 2010; Long et al. 2016; Panigrahi and O’Malley 2021). In *Drosophila*, a few *cis*-regulatory regions (non-coding regions flanking coding DNA) have been dissected in detail (Small and Arnosti 2020). These analyses have isolated compact elements that can largely recapitulate, in whole or part, the endogenous expression pattern of the target gene(s). For example, the pattern of seven stripes of the pair-rule gene *even*-*skipped* (*eve*) in the early embryo of *D. melanogaster* is determined by several small “minimal” enhancers, each directing expression in one or two stripes (Fujioka et al. 1999; Sackerson et al. 1999). Results such as these imply that the regulatory information that constitutes an enhancer is confined to a compact DNA region with autonomous function.

In contrast, several lines of evidence suggest that minimal enhancers do not include all the regulatory information for driving gene expression. For example, despite the early work defining minimal enhancers of the *eve* locus, a large reporter construct of the *eve* locus with a deletion of the minimal stripe 2 element (the enhancer that generates the second of the seven stripes) retains residual expression at stripe 2 cells (Ludwig et al. 2005). In addition, it was shown that regions outside of the stripe 2 minimal enhancer buffer gene expression under environmental or genetic perturbations (Ludwig et al. 2011; López-Rivera et al. 2020). Furthermore, in *D. erecta*, a species closely related to *D. melanogaster,* TFBSs located outside of the minimal stripe 2 enhancer region are needed for expression of *eve* in stripe 2 (Crocker and Stern 2017). Other evidence also calls into question the ubiquity of compact regulatory regions in *Drosophila*. In the *D. melanogaster* early embryo, six of the seven stripes of the pair-rule gene *hairy* are generated by seemingly compact enhancers, but regulatory information for driving *hairy* stripe 2 is spread over many kilobases (Riddihough and Ish-Horowicz 1991). Similarly, several aspects of the expression of the pair-rule gene *runt* in *D. melanogaster* appear to be mediated by regulatory information scattered over many kilobases (Klingler et al. 1996). The notion that regulatory information may be spread over kilobases linked to the possibility that a coherent expression pattern might only emerge upon interaction of scattered sub-elements (Frankel 2012) could explain the inability to isolate discrete enhancers from large regulatory regions in particular cases (Davis et al. 2007).

Evolutionary analyses have estimated that the fraction of functionally relevant nucleotides (nucleotides that evolve under either negative or positive selection, and thus have a function) in intergenic regions of the *D. melanogaster* genome is ∼0.6 (Andolfatto 2005; Halligan and Keightley 2006). This estimate suggests that most nucleotides in *cis*-regulatory regions are functional and are potentially involved in the control of gene expression. But, how can we explain such a large fraction of functional nucleotides if regulatory elements occupy small segments of non-coding DNA? How much of the non-coding constraint within the *Drosophila* genome is explained by the presence of regulatory DNA?

Here, we explore this issue by examining the distribution of conserved elements and the function of non-coding DNA in the regulatory region of the *shavenbaby* (*svb*) gene. Svb is a transcription factor that controls the formation of non-sensory cuticular hairs (trichomes) in the larva (Payre et al. 1999) and pupa (Delon et al. 2003; Preger-Ben Noon, Sabarís et al. 2018) of *D. melanogaster*. The *svb* regulatory region has been scrutinized for decades, providing a solid platform for exploring mechanistic and evolutionary aspects of gene regulation (Frankel et al. 2012; Stern and Frankel 2013; Kittelmann et al. 2021; Soverna et al. 2021). Prior comprehensive analyses of the ∼90 kb region upstream of the *svb* first exon using reporter constructs revealed that regulatory activity in the embryo is limited to seven enhancers, some of which have been dissected to fragments of less than 1 kilobase (McGregor et al. 2007; Frankel et al. 2010; Frankel et al. 2011; Crocker et al. 2015; Preger-Ben Noon et al. 2016; Preger-Ben Noon, Sabarís et al. 2018). These seven enhancers are also active in the epidermis of the pupal abdomen and other larval tissues (Preger-Ben Noon, Sabarís et al. 2018).

Here, we verified that regions outside of the known enhancer elements display high sequence conservation (Stern and Frankel 2013). We reasoned that there might be additional regulatory information in the locus controlling *svb* expression in contexts other than the embryo. Thus, we undertook a functional characterization of the *cis*-regulatory region of *svb* in the epidermis of the pupal abdomen. As opposed to achieving *svb* activation through compact and disperse enhancers, we found that *svb* expression in the abdominal pupal epidermis results from the activity of large DNA regions with enhancer activity, which occupy most of the ∼90 kb upstream of *svb* first exon.

To assess the generality of our finding, we scrutinized regulatory regions of other developmental genes. We observed that other regulatory regions of developmental genes also display dense collections of putative regulatory elements and that a large fraction of conserved bases lie within these putative regulatory elements. We hypothesized that this pattern might be extrapolated to the whole genome and therefore performed a genome-wide analysis. We found that a large fraction of conserved non-coding DNA throughout the genome is contained within putative regulatory DNA. Overall, these results suggest that the widespread conservation of *Drosophila* non-coding DNA can be explained, at least in part, by the distribution of regulatory elements throughout the non-coding genome.

## Results

### The regulatory region of the svb gene displays widespread sequence conservation

We analyzed sequence conservation patterns of the *svb cis*-regulatory region and other non-coding genomic regions of similar size using multi-species genome alignments (see Materials and Methods for details). First, we calculated phastCons values (Siepel et al. 2005) throughout the *svb cis*-regulatory region (Figure 1A). We observed similar fractions of bases in phastCons conserved elements in the seven known enhancers and the regions that lie outside enhancers (Figure 1B, Table S1). Second, we found that the fraction of bases in phastCons conserved elements in the whole *cis*-regulatory region of *svb* is greater than 80% of 10,000 randomly chosen windows of non-coding DNA of the same size in the *D. melanogaster* genome (Figure 1C). We reasoned that the high sequence conservation could be indicative of the existence of functional non-coding DNA, so we embarked on a search for additional regulatory elements. Since we had identified all enhancers that drive *svb* expression in the embryonic epidermis (McGregor et al. 2007; Frankel et al. 2010), we decided to characterize *svb* expression in the pupal abdominal epidermis, another tissue in which Svb is required for trichome production (Preger-Ben Noon, Sabarís et al. 2018).

**Figure 1.**
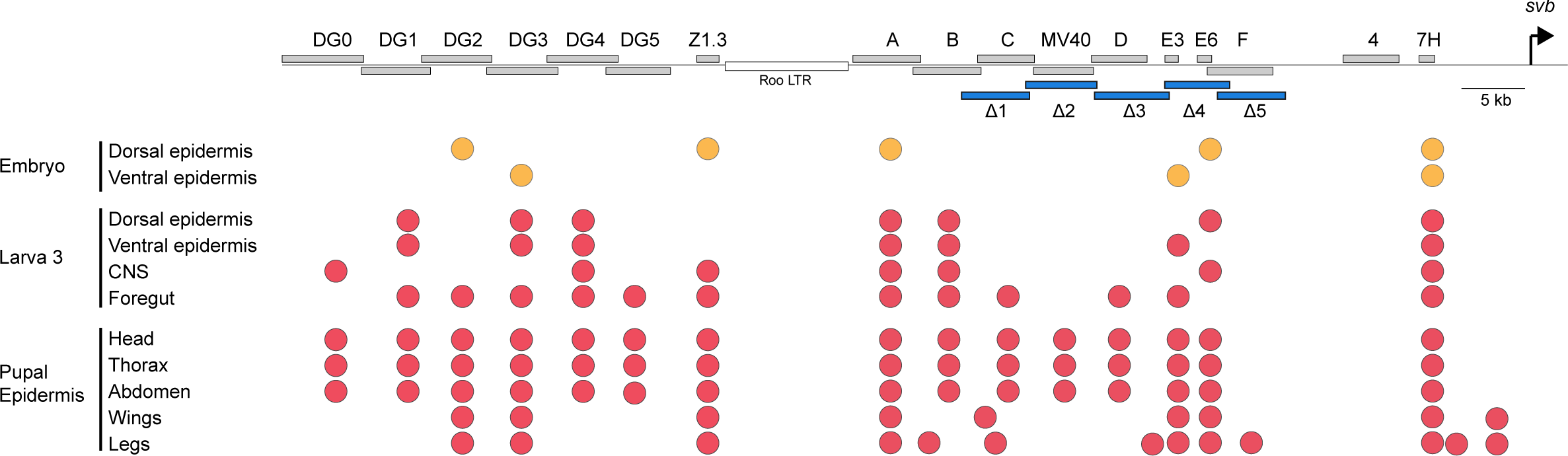
Widespread sequence conservation in the regulatory region of the *svb* gene. (A) Scheme of the *svb cis*-regulatory region of *D. melanogaster*, showing the position of the embryonic enhancers (grey boxes) and the pattern of sequence conservation in a multiple sequence alignment of 23 *Drosophila* species and 4 outgroup species. Orange peaks represent phastCons conservation scores per base. Blue boxes represent conserved elements predicted by phastCons. In *D. melanogaster*, the regulatory region of *svb* contains a species-specific transposable element (Roo LTR) (B) Boxplots with the fraction of bases in conserved elements in the seven enhancers (left) and eight regions that lie outside enhancers (right). Empty circles indicate values for each DNA fragment. The differences between categories are not significant (Mann-Whitney test, p=0.61) (C) Density plot showing sequence conservation in different non-coding regions of the *D. melanogaster* genome. The x-axis indicates the fraction of bases within phastCons conserved elements in 10000 non-coding windows of the *D. melanogaster* genome. The red dashed line indicates the fraction of conserved bases for the *svb cis*-regulatory region (0.511).

### Svb is active in the abdominal epidermis of pupae at ∼40-45 hours after puparium formation

To investigate *svb* expression in the pupal epidermis we examined expression driven by a BAC carrying the regulatory region of *svb* upstream of a GFP-NLS reporter, named *svb*BAC-GFP (Figure 2A; Preger-Ben Noon, Sabarís et al. 2018). We observed GFP in larval epidermal cells at 20 hours after puparium formation (hAPF), which is consistent with earlier *svb* expression linked to the formation of the puparium (Figure 2A). The GFP signal observed in larval epidermal cells disappears during histolysis of these cells (Figure 2A). The pupal abdominal epidermis is derived from histoblast nests cells, which divide and migrate in the early pupa, replacing the larval epidermis across the whole abdomen. We first detected GFP expression in pupal epidermal cells at 35 hAPF (Figure 2A). Later on, GFP levels increase, and by 45 hAPF all pupal abdominal epidermal cells display bright GFP signal.

**Figure 2.**
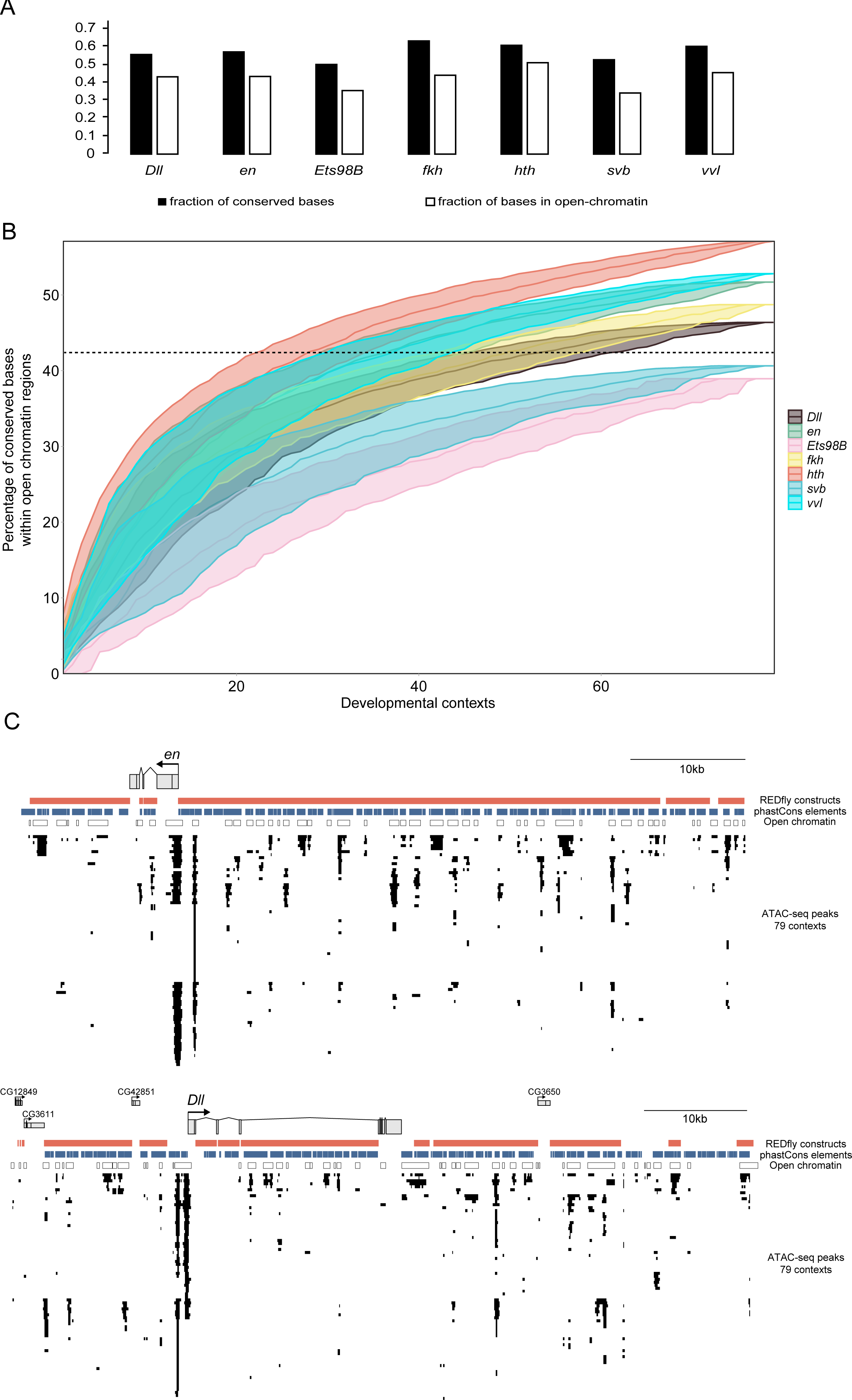
*svb* expression and Svb activation in the abdominal pupal epidermis of *D. melanogaster* (A) GFP expression driven by the *svbBAC*-GFP reporter construct (scheme above) in larval epidermal cells (large nuclei) and pupal epidermal cells (small nuclei) of the pupal abdomen between 20 and 45 hours APF (B) Presence of the full length Svb (transcriptional repressor) in nuclei of the abdominal pupal epidermis between 39 and 45 hours APF (magenta). Images below show the DAPI signal (blue) for the same fields.

To determine whether the patterns of GFP transcription driven by the *svb*BAC-GFP reflect the expression of Svb protein, we stained abdominal epidermis with an antibody that recognizes the N-terminus of Svb (Chanut-Delalande et al. 2014). Full-length Svb protein acts as a transcriptional repressor and is converted into a transcriptional activator upon degradation of its N-terminus (Kondo et al. 2010). We detected Svb repressor in the epidermis of the pupal abdomen at 39 hAPF (Figure 2B). Starting at approximately 41 APF, the intensity of Svb repressor staining decreases, until it becomes almost undetectable at 45 hAPF (Figure 2B). These results suggest that Svb is converted into a transcriptional activator between 40 and 45 hAPF, which implies that Svb target genes are activated during this interval, triggering trichome development. Our data are consistent with a previous study that identified the beginning of trichome formation in the abdominal epidermis at approximately 45 hAPF (Mangione and Martín-Blanco 2018).

### Chromatin landscapes of the svb regulatory region in the embryo and pupa are sharply different

To characterize the regulatory landscapes of *svb* in the epidermis of the embryo and pupal abdomen we assayed open chromatin and the presence of the histone mark H3K27ac, which together can provide substantial evidence of the existence of active enhancers (Moore et al. 2020). We performed fluorescently activated cell sorting (FACS) to isolate *svb*-expressing epidermal cells from late embryos and performed chromatin immunoprecipitation (ChIP)-seq to quantify the genome-wide H3K27ac signal in these cells. In addition, we characterized open chromatin regions by retrieving computationally-defined clusters of single-cell ATAC-seq data corresponding to epidermal cells of the late embryo (Cusanovich et al. 2018). We analyzed these data in the context of previous findings, which uncovered that seven enhancers scattered in the *svb* regulatory region generate *svb* expression in the embryo (in previous experiments the whole ∼90 kb upstream of *svb* were scrutinized for embryonic enhancers through reporter constructs). We observed that genomic regions showing high levels of open chromatin and H3K27ac enrichment are largely coincident with the locations of the seven previously characterized embryonic enhancers (Figure 3). In fact, there is a consistent overlap of ATAC peaks and flanking acetylation signals specifically for enhancer regions that have been dissected to “minimal elements” (Z1.3, E3, E6 and 7H, Figure 3). Furthermore, ATAC-seq peaks and acetylation signal within enhancers that have not been dissected to small elements (DG2, DG3 and A) are potential predictors of the position of “minimal elements” (Figure 3). Thus, the chromatin landscape of the *svb* locus in epidermal cells of the embryo reveals that embryonic enhancers are small islands in a large regulatory region, which is consistent with prior knowledge from reporter assays.

**Figure 3.**
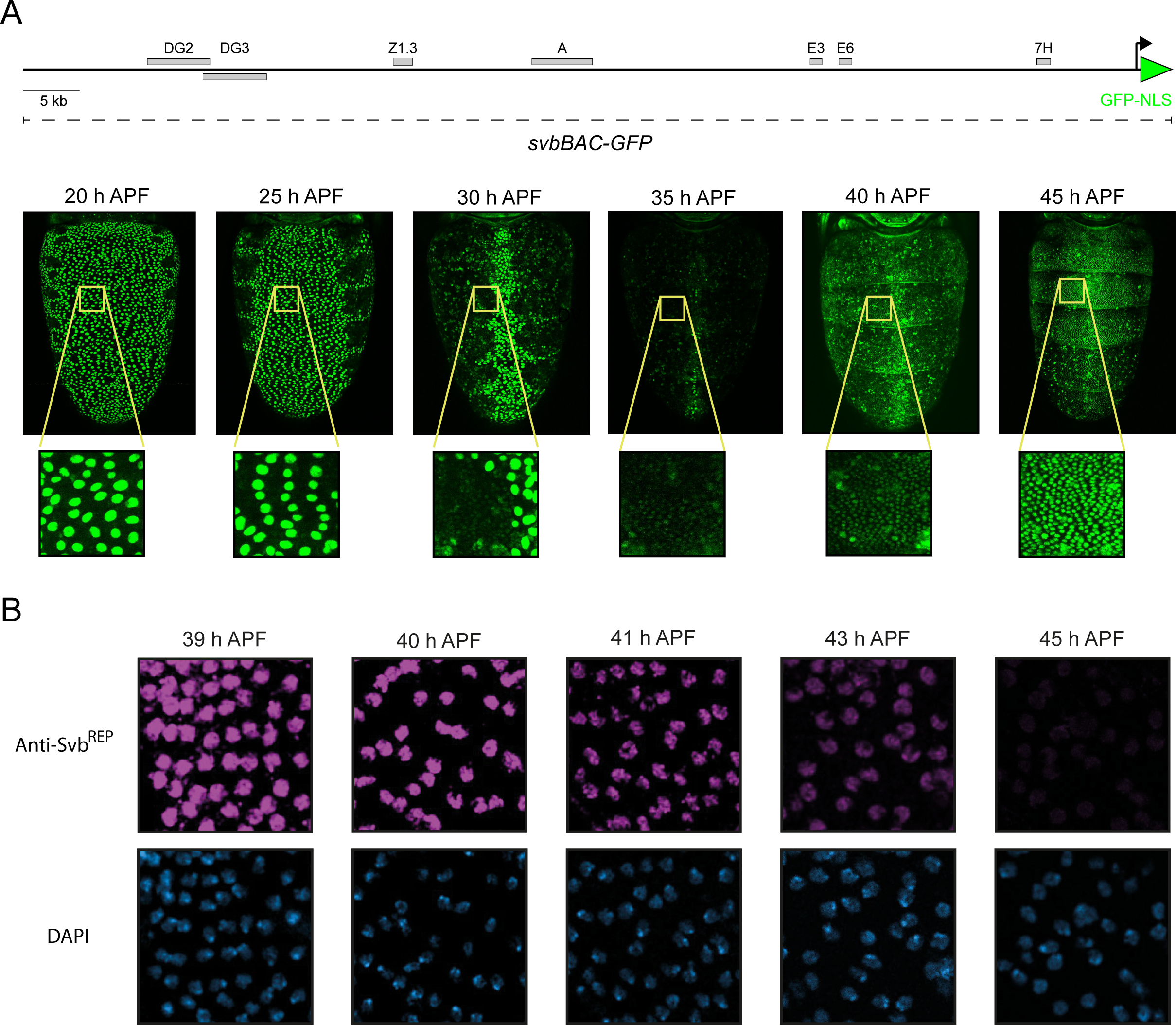
Contrasting chromatin landscapes in the *svb* regulatory region between embryonic and pupal epidermis. Chromatin architecture of the *svb* locus in the embryonic epidermis (above) and the pupal abdominal epidermis (below). ATAC-seq profiles are colored in blue and ATAC-seq peaks are indicated with black boxes below profiles. H3K27ac profiles are colored in red. Embryonic enhancers of *svb* are represented with grey boxes. Gray shading marks the position of embryonic enhancers in embryonic tracks. In pupal tracks, gray shading marks the location of regions with high density of ATAC-seq peaks and H3K27ac signal.

To characterize the chromatin landscape of the *svb* locus in epidermal cells of the pupal abdomen, we dissected the epidermis of the pupal abdomen between 38-45 hAPF and performed ATAC-seq and CUT&RUN against the H3K27ac mark. Strikingly, we observed large areas of the *svb* upstream region displaying ATAC-seq peaks and elevated H3K27ac signal (Figure 3). Notably, most of these regions are not coincident with minimal embryonic enhancers, but rather correspond to DNA regions that lie between the embryonic enhancers (Figure 3). Thus, the chromatin landscapes of the *svb* regulatory region in epidermal cells of the embryo and epidermal cells of the pupa are remarkably different (Figure 3). Altogether, these results suggest that regulatory DNA driving pupal epidermal expression is spread across large regions that are not active in the embryo. Furthermore, open chromatin data from other tissues in which *svb* is expressed suggest that the regulatory landscape of this gene might be highly variable between contexts (Supplementary Figure 1). Given that chromatin structure in epidermal cells of the pupal abdomen suggests that the *svb cis*-regulatory region contains multiple enhancer elements, we sought to further validate these candidate enhancer regions with reporter constructs.

### Reporter gene assays validate novel pupal and larval enhancers of the svb gene

Previously, we showed that the seven enhancers with embryonic activity are also active in the pupal epidermis (Preger-Ben Noon, Sabarís et al. 2018). Here, we examined reporter constructs that encompass most of the remaining *svb* upstream region (Figure 4). We analyzed the activity of these fragments, which are cloned upstream of the *lacZ* reporter gene, in whole pupae. We observed that all eight fragments within candidate enhancer regions in the pupa (fragments DG0, DG1, DG4, DG5, B, C, D and MV40) drove reporter expression in the pupal abdominal epidermis (Figure 4, Supplementary Figure 2). All of these fragments are also active in the epidermis of the thorax and head (Figure 4, Supplementary Figure 2). Two fragments that lie outside of the candidate enhancer regions (fragments F and 4) did not display enhancer activity in the pupal epidermis (Figure 4, Supplementary Figure 2). Altogether, these analyses confirm that multiple DNA regions encode regulatory information capable of driving reporter expression in the pupal epidermis.

**Figure 4.**
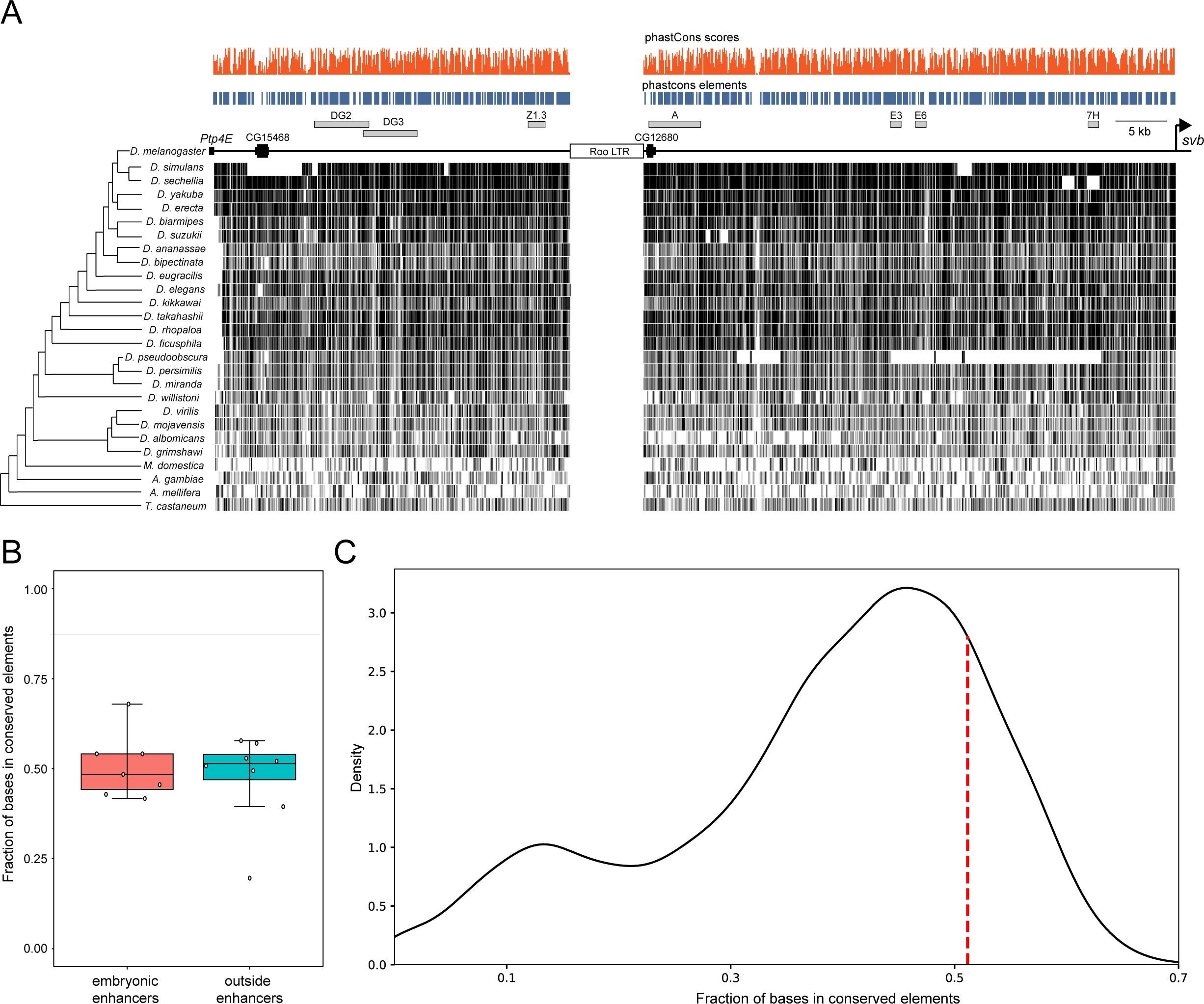
Reporter constructs within the candidate regulatory regions have enhancer activity in pupa. Schematic of expression patterns driven by *svb* reporter constructs in the embryo and pupal epidermis (the panel integrates expression data obtained in McGregor et al (2007), Frankel et al (2010), Preger-Ben Noon, Sabarís et al (2018) and this study). The diagram of the *svb* locus shows the position of the seven enhancers that are active in the embryo, larva and pupa (gray boxes) and ten fragments without enhancer activity in the embryo (orange boxes) that were examined for pupal enhancer activity in this study. The position of candidate regulatory regions is indicated with light-gray boxes.

In an earlier study, we demonstrated that the original seven *svb* enhancers (DG2, DG3, A, Z1.3, E3, E6 and 7H) drive expression in the embryo, larva and pupa (Preger-Ben Noon, Sabarís et al. 2018) and wondered whether the newly characterized fragments with pupal expression may also be active in other developmental contexts. Preliminary evidence from reporter constructs suggested that *svb* is expressed in the brain, imaginal discs and foregut of the larva (Preger-Ben Noon, Sabarís et al. 2018), but the presence of the protein in these organs had not been confirmed. We used the antibody against the N-terminus of the Svb protein for immunofluorescence assays and confirmed that Svb is indeed present in brain, imaginal discs and foregut of the larva (Supplementary Figure 3). We tested the activity of reporter constructs in these organs and in the epidermis of third instar larvae (Supplementary Figure 4). We observed that seven of the eight constructs that had pupal activity also drove expression in at least one larval organ (Supplementary Figure 4). Six constructs drove expression in the foregut, 3 in the epidermis, and 3 in the brain. Surprisingly, none of these reporter constructs drove expression in imaginal discs, even though some of these regions appear to have open chromatin in this organ (Supplementary Figure 1)

### Novel pupal enhancers are required for svb expression

Reporter constructs provide evidence that multiple DNA fragments within the *svb cis*-regulatory region contain information for driving the wild type *svb* expression pattern. To determine whether these regions are required for *in vivo* activity, we used BAC recombineering to generate five mutant versions of *svb*BAC-GFP, each containing a deletion of approximately 5 kb. Four deletions were made in regions with enhancer activity in reporter assays (Δ1-Δ4) and one deletion was made in a region with no enhancer activity (Δ5) (Figure 5A). These five versions of *svb*BAC-GFP were integrated into a specific attP site of the *D. melanogaster* genome. To normalize the fluorescence signal we compared the GFP signal from mutant *svb*BACs with the DsRed signal from a wild-type *svb*BAC (*svb*BAC-DsRed) integrated into a different attP site, to avoid transvection (Mellert and Truman 2012). We quantified expression of the GFP BACs carrying deletions and *svb*BAC-DsRed in the epidermis of the dorsal abdomen (Figure 5B). Two deletions, both from regions with enhancer activity in reporter assays, had effects on GFP expression (Δ3 and Δ4). Remarkably, while Δ3 reduced expression level, deletion of a neighboring region (Δ4) increased expression (Figure 5B). The increase in GFP expression in Δ4 was also observed when deleting the pleiotropic enhancer E6 in *svb*BAC-GFP, which is included in the fragment deleted in Δ4 (Preger-Ben Noon, Sabarís et al. 2018). Δ1 and Δ2 diminished mean expression levels slightly but not significantly (Figure 5B). The deletion of a region with no enhancer activity (Δ5) had no significant effects on GFP expression (Figure 5B).

**Figure 5.**
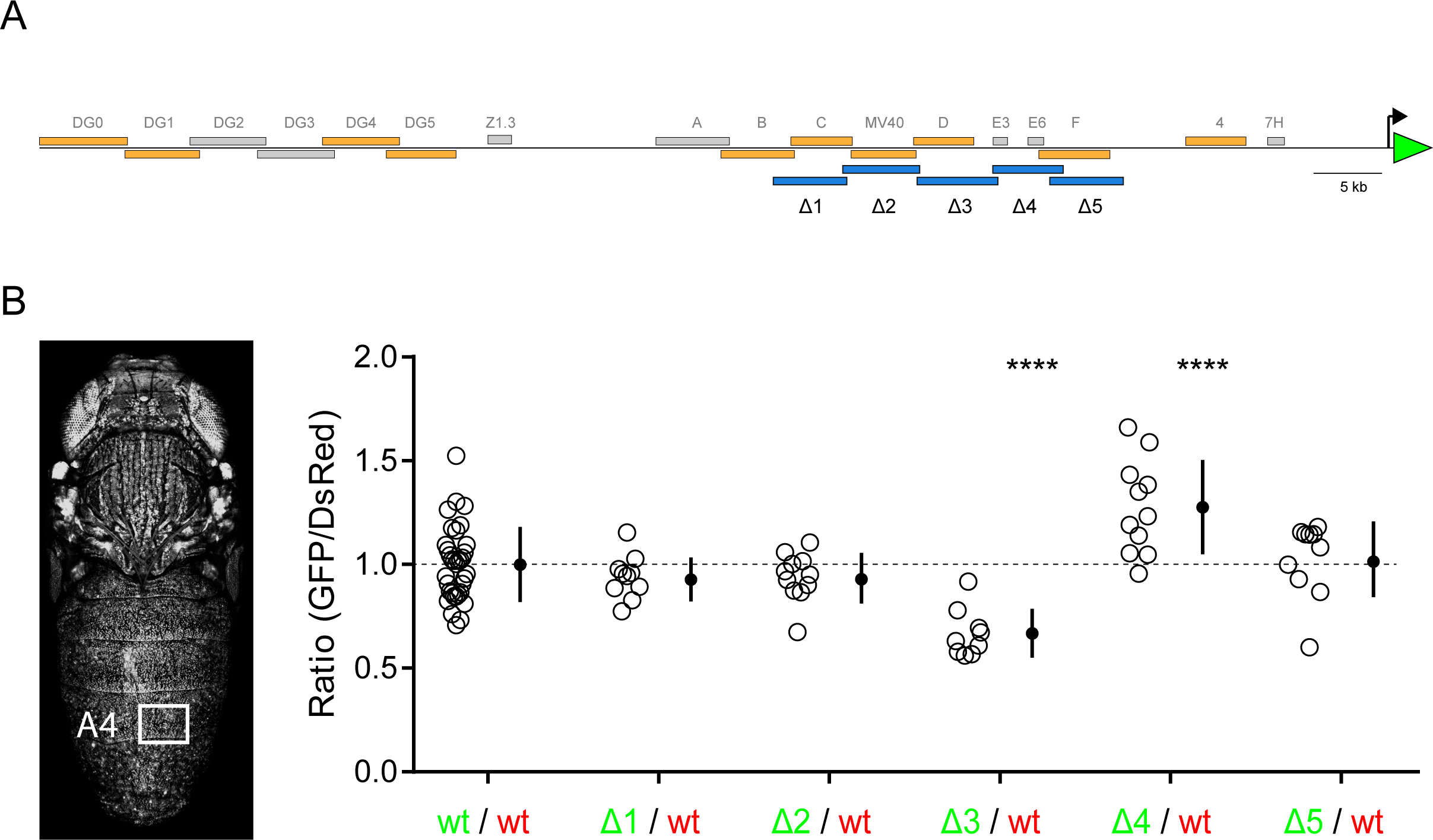
DNA within one of the large regions with presumptive regulatory activity bears pupal enhancer activity *in vivo*. (A) Position of deletions (blue boxes) that were generated in a BAC reporter construct carrying the regulatory region of *svb* and a GFP reporter (*svb*BAC-GFP). A wild type *svb*BAC-DsRed BAC was used to normalize GFP levels (B) Effect of deletions on pupal expression. The GFP/DsRed ratio was measured in part of abdominal segment A4 (rectangle). Open circles indicate the average ratio (GFP/DsRed) for each individual. Closed black circles and vertical lines indicate mean and 1 SD, respectively. The dashed line marks the mean ratio of wild type constructs. Statistical significance was calculated using one-way ANOVA and Dunnett’s pairwise comparisons (**** p<0.0001).

These results reveal, first, that DNA sequences outside of previously characterized enhancers are required for wild type *svb* expression in the pupal epidermis. Second, deletions of two regions with enhancer activity in reporter assays do not alter gene expression, suggesting that their activity is buffered by other DNA regions with similar expression patterns, as has been observed for *svb* embryonic expression (Frankel et al. 2010). Finally, the increase in expression in Δ4 may be explained with a model of enhancer-promoter competition (Bothma et al. 2015) or, alternatively, with a scenario in which Δ4 contains regulatory elements that both enhance and silence gene expression. Altogether, these findings suggest that, despite the observed redundancy in the activity of *lacZ* constructs, DNA pieces within *svb* regulatory region may not have an equivalent function *in vivo*.

### Regulatory function appears as a major determinant of non-coding sequence conservation in the svb locus and throughout the Drosophila genome

We built an atlas of regulatory elements for the *svb* locus (Figure 6) using data from this work along with information collected in our previous studies in embryo, larva and pupa (McGregor et al. 2007; Frankel et al. 2010; Preger-Ben Noon, Sabaris et al. 2018; Kittelmann et al. 2018). This atlas shows, prominently, that regulatory information for driving *svb* expression is scattered throughout the whole *cis*-regulatory region (Figure 6). Embryonic enhancers do not occupy much of the *cis*-regulatory region. However, if we also consider *svb* expression in different tissues of the larva and pupa, the regulatory elements associated with these contexts cover most of this non-coding DNA (Figure 6). This regulatory architecture is a likely cause of much of the sequence conservation throughout the locus.

**Figure 6.**
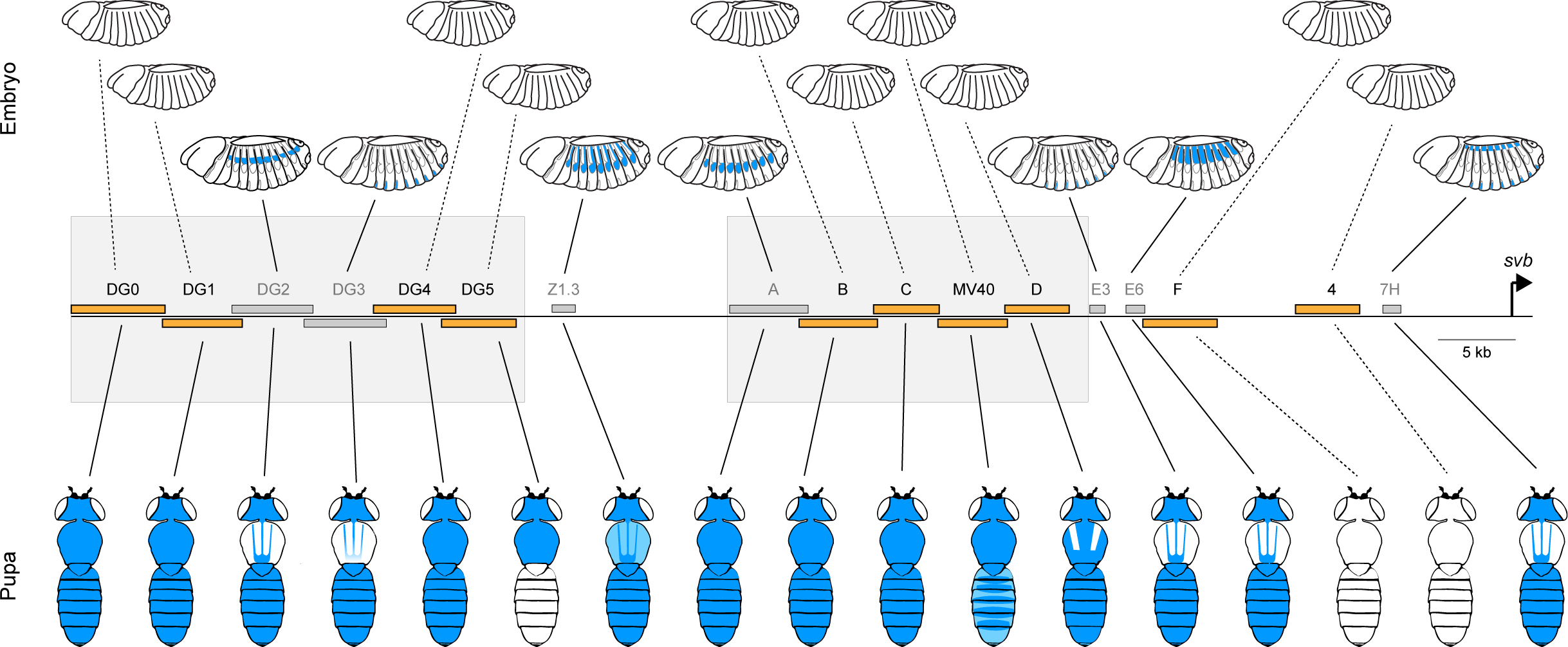
A myriad of enhancers in a large regulatory region. Summary of enhancer activities in the *svb* locus in embryo, larva and pupa. Orange circles indicate embryonic activity and red circles indicate larval/pupal activity. Expression data derives from previous works (McGregor et al. (2007), Frankel et al. (2010), Kittelmann et al. (2018), Preger-Ben Noon, Sabarís et al. (2018)) and this study.

We wondered if this link between the density of regulatory and the extent of sequence conservation is a general phenomenon. Hence, we decided to explore a possible relationship between non-coding sequence conservation and the abundance of regulatory information in other developmental genes and throughout the *Drosophila* genome. To determine whether sequence conservation can be explained by the density of regulatory DNA, we intersected ATAC-seq peaks for 79 developmental contexts with the collection of phastCons non-coding conserved elements of the *Drosophila* genome. We first inspected the regulatory regions of well-studied developmental genes that possess large non-coding regions in *cis* (Nelson et al. 2004) and that are known to be active across multiple developmental contexts (*Dll*, *en*, *Ets98B*, *fkh*, *hth*, *vvl* and *svb;* Figure 7). These large regulatory regions were defined by the position of validated regulatory elements (see Figure 7C). We observed that these regulatory regions have a high density of putative regulatory elements, since 34% to 50% of their bases lie within open chromatin when summing all 79 developmental contexts (Figure 7A). As well, we calculated that between 49% and 63% of their bases are found in conserved elements (Figure 7A). With these data, we examined the fraction of conserved bases that fall within ATAC-seq peaks. We calculated how this percentage changes with the number of contexts that are considered (Figure 7B). We observed that this parameter first grows rapidly, and that the slope decreases when about 20 contexts have been included in the analysis. However, curves do not seem to reach an asymptote (Figure 7B). When considering all contexts, 39% to 57% of the conserved bases in these regulatory regions fall within open chromatin (Figure 7B). To rule out that the overlap between open chromatin and conserved bases is a mere result of the abundance of these two features in the genome, we performed an odds ratio analysis with the number of conserved and non-conserved bases that occur outside open-chromatin peaks. Since regulatory elements can evolve rapidly (Swanson et al. 2010), not all their bases are expected to be conserved. However, an odds ratio may indicate whether there is an enrichment of conserved bases in putative regulatory elements (open-chromatin regions). Indeed, we observed that conserved bases are more likely to be found in open chromatin than non-conserved bases (for all regulatory regions odds ratios are significantly higher than 1, Fisher’s exact test p<0.00001, see Figure S5A).

**Figure 7.**
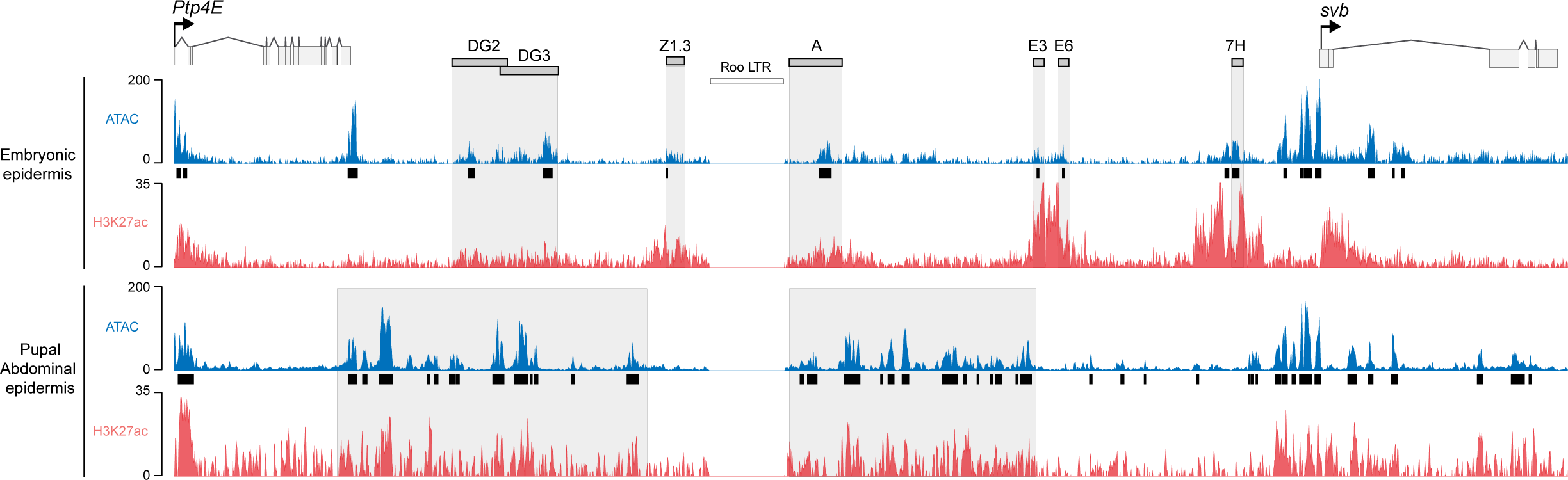
Extensive overlap between putative regulatory elements and phastCons elements in regulatory regions of developmental genes and throughout the non-coding *Drosophila* genome (A) Fraction of conserved bases (black bars) and fraction of bases in open-chromatin considering all developmental contexts (white bars) in regulatory regions of developmental genes (B) Overlap between phastCons elements and open chromatin in regulatory regions of developmental genes considering different number of developmental contexts. Curves vary slightly depending on the order of sampling of contexts. Colored lines indicate average values. Shaded bounding areas indicate upper and lower values. The dotted line indicates the average overlap for 10,000 random windows, considering all 79 contexts (C) Overlap between conservation and open chromatin in the regulatory regions of the developmental genes *engrailed* (above) and *Distal-less* (below). Black boxes indicate regions of open chromatin for each of the 79 develomental contexts. White boxes indicate regions of open chromatin for all 79 develomental contexts. *phastCons* elements are marked in blue. Red boxes indicate regions that are part of reporter constructs with validated regulatory activity (REDFly constructs). These data were used to generate the graph of panel A.

We also calculated the percentage of bases in conserved elements that are within ATAC-seq peaks in the 10,000 random windows of the genome that we had generated for the analysis of Figure 1C (these windows have the size of the regulatory region of *svb*). We considered all 79 developmental contexts for this calculation. We found that, on average, 42.4% of the bases in conserved elements are within putative regulatory DNA (dotted line in Figure 7A). This average is quite similar to the value obtained for the *svb* regulatory region counting all 79 developmental contexts (40.6%). In 82.5% of these 10,000 windows, we also observed that conserved bases are more likely to be found in open chromatin than non-conserved bases (odds ratios are higher than 1 in 82.5% of the windows, see Figure S5B)

## Discussion

At many loci in the *Drosophila* genome, non-coding regions display high levels of sequence conservation that cannot be explained by the density and distribution of known *cis*-regulatory elements. This is clearly demonstrated at the *svb* locus, where the seven embryonic enhancers that we identified previously (McGregor et al. 2007; Frankel et al. 2010) cannot account for the high level of sequence conservation. In this work, we showed that the density of regulatory information can explain, at least in part, sequence conservation in the regulatory region of the *svb* gene Through functional analyses we demonstrated that the regulatory landscapes of the *svb* locus in embryo and pupa are remarkably different. While the activation of *svb* in the embryo is driven by small and discrete enhancers, pupal *svb* expression is generated by tens of kilobases of almost continuous regulatory information. Thus, the expression of a single transcriptional factor, which very likely activates the same target genes for producing trichomes in embryonic and pupal epidermis, is regulated through completely different mechanisms in the two contexts.

Close examination of trichome morphology reveals that sizes and shapes of trichomes vary over the embryo, larva and adult, and previous work has shown that Svb levels modulate trichome number and size (Delon et al. 2003). Thus, it is conceivable that different levels of *svb* are needed in the embryo and pupa to generate the right number of trichomes with the right size, and that the seemingly redundant expression patterns driven by multiple regulatory elements are required to define precise quantitative levels of transcription at different times and in different places. It is possible that in the pupal epidermis the whole 90 kb *cis*-region of *svb* acts as a single regulatory unit to ensure the correct transcriptional output for the *svb* gene. Such a feature has been observed during mouse embryogenesis, where multiple redundant enhancer elements act in concert to activate the genes *Ihh* (Will et al. 2017) and *Fgf8* (Hörnblad et al. 2021). Similarly, the transcriptional activity of Locus Control Regions (LCRs) and so-called “super-enhancers” is achieved through the action of multiple small enhancer elements (Grosveld et al. 2021).

We explored whether the pattern of conservation and density of regulatory elements observed in the *svb* locus is reproduced in other developmental genes and throughout the *Drosophila* genome. We found that other regions also have a high density of putative regulatory elements. By analyzing the overlap between conserved elements and open chromatin we showed that more than 40% of the bases in conserved elements of the *Drosophila* genome fall within putative regulatory elements. Furthermore, in some regulatory regions of developmental genes this percentage rises above 50%. This finding begs the question as to whether conserved DNA can be explained just by the presence of regulatory DNA. To get an answer for this question it will be necessary to obtain ATAC-seq data for many more developmental contexts and to further validate the function of putative regulatory elements throughout the genome.

The density of putative regulatory activity that we detected at *Drosophila* developmental genes appears different from that of vertebrate developmental genes. For example, enhancers of the *HoxD* cluster and *Shh* and *Pax6* genes in mice are separated by tens of kilobases of non-coding DNA that do not seem to encode a function (Montavon et al. 2011; Anderson et al. 2014; Buckle et al. 2018). Similarly, patterns of phylogenetic footprinting in non-coding DNA of vertebrates indicate that conserved regions are separated by large non-conserved sequences (Santini et al. 2003; Sun et al. 2006; Navratilova et al. 2009; Peterson et al. 2009). In contrast to vertebrates, the high density of regulatory information in non-coding DNA of *Drosophila* may simply result from the compactness of its genome (Nelson et al. 2004).

Altogether, our results suggest that the non-coding *Drosophila* genome is dense in regulatory information. Thus, the widespread understanding of regulatory information as encoded in compact and dispersed elements, which has been implied by studies of regulatory regions mostly in the *Drosophila* embryo, may not be representative of the architecture of regulatory information. A future challenge is to determine how this plethora of regulatory information is integrated in space and time to achieve precise regulatory outputs.

## Materials and methods

### Multiple sequence alignment of the svb cis-regulatory region and quantification of evolutionary conservation

We defined the *svb* regulatory region in *Drosophila melanogaster* as the segment between the *svb* TSS and the last base of the coding region of the gene *Ptp4E* (92355 bp). Coordinates of the *svb* regulatory region in most species were obtained from an existing multi-species genome alignment (27-way alignment in the USCS Genome Browser). We filtered the MAF file with MafFilter (Dutheil et al. 2014) to keep the sequences orthologous to the *svb* regulatory region of *D. melanogaster*. Since we could not find the whole regulatory region of *svb* for *D. persimilis*, *D. mojavensis, M. domestica, A. mellifera, A. gambiae and T. castaneum* in the alignment file, we searched for the coordinates of the complete region using BLASTP with the protein sequences of Ptp4E and Svb as queries (these two coding regions flank the regulatory region of *svb* in *D. melanogaster*). For the non-*Drosophila* species we observed that *Ptp4E* was > 200 kb away from *svb* first exon, so we defined the *svb* regulatory region in these species as the 150 kb sequence upstream of *svb* TSS. We extracted the sequences of the *svb* regulatory region from all 26 species with Bedtools (Quinlan and Hall 2010). We generated a multiple alignment of the *svb* regulatory region using the local multiple sequence aligner TBA from multiz package (Blanchette et al. 2004). To perform the multi-alignment, TBA generates a series of pair-wise alignments to “seed” the multiple alignment process. We performed the pair-wise alignments using an optional “blastz specs file” with the following parameters: --hspthresh=1500 – gappedthresh=1500. Measurements of evolutionary conservation in the *svb* regulatory region were performed with PhastCons from the PHAST package (Siepel et al. 2005) with the following parameters: expected-length=45, target-coverage=0.3, rho=0.3. The non-conserved model was downloaded from UCSC site (https://hgdownload.soe.ucsc.edu/goldenPath/dm6/phastCons27way/). To identify conserved elements, we ran PhastCons with the --most-conserved --score parameters. We removed poorly conserved elements (elements < 25 bp and with log odds scores < 60). Tracks were visualized using the UCSC Genome Browser. Genome-wide measurements of evolutionary conservation were performed using PhastCons data based on the existing multi-species genome alignment from UCSC (phastCons27way track). Again, we removed poorly conserved elements (elements < 25 bp and with log odds scores < 60). We removed all exons and transposons not overlapping phastCons elements from the alignment. 10000 random windows of 82184 bp (the length of the *svb* regulatory region after removing exons and the Roo transposable element) were selected from the *D. melanogaster* genome. The fraction of conserved bases (bases within conserved elements/total number of bases) was calculated using Bedmap in BEDOPS suite (Neph et al. 2012) with parameters --echo --delim ‘\t’ –-bases-uniq, with the windows bed file as reference file and the conserved elements bed file as map file.

### Overlap between conserved elements and open chromatin in developmental genes and throughout the genome

To study the relationship between sequence conservation and regulatory activity in non-coding regions of the genome we downloaded processed bulk ATAC-seq peaks from ChIP-ATLAS (https://chip-atlas.org/) and single-cell ATAC-seq embryonic data from scEnhancer (http://enhanceratlas.net/scenhancer). We also included ATAC-seq peaks from abdominal pupal epidermis generated in this work. We merged peaks from the same developmental context when more than one source was available. The whole dataset consisted of 79 distinct contexts, including embryonic cell types, and larval, pupal and adult tissues (Supplementary Table 3).

We defined regulatory regions for seven classical developmental genes (*Dll, en, Ets98B, fkh, hth, svb and vvl*) by using REDfly (redfly.ccr.buffalo.edu). For each gene, the regulatory region was defined as the DNA region between the two most distant validated regulatory elements for the gene. We calculated the overlap between conserved bases (bases in conserved phastCons elements) and open chromatin regions. 500 growth curves were generated for each regulatory region by sampling an increasing number of developmental contexts (from all 79 contexts) in different orders.

For obtaining odds ratios we first calculated the number of conserved bases within (A) and outside (B) open chromatin (odds = A/B). Next, we calculated the number of non-conserved bases within (C) and outside (D) open chromatin (odds = C/D). The odds ratio for each regulatory region = AD/BC. We performed the same calculation for the 10,000 random windows. We excluded 61 random windows because they had zero conserved or non-conserved bases. For the seven regulatory regions, we used Fisher’s exact tests to determine the statistical significance of the enrichment of conserved bases in open chromatin.

### Fly strains

Enhancer-*lacZ* reporter lines B, C, MV40, D, F and 4 are described in McGregor et al (2007), while enhancer-*lacZ* reporter lines DG0, DG1, DG4 and DG5 are described in Frankel et al (2010). *svb*BAC-GFP and *svb*BAC-DsRed lines are described in Preger-Ben Noon, Sabarís et al (2018). We used BAC recombineering (Wang et al. 2009) to delete regions of approximately 5 kb in the context of the *svb*BAC-GFP. All primers and constructs that were used for BAC recombineering are listed in Table S2. These constructs were integrated into the fly genome through attP/attB recombination (Rainbow Transgenic Flies Inc.). The different versions of *svb*BAC-GFP were integrated in attP site VK00033. *svb*BAC-DsRed was integrated in attP site VK00037.

### Immunofluorescence in pupa and larva

Larva 3 tissues were dissected, fixed and stained using a standard protocol with anti-Svb1s (1:300) and anti-rabbit Alexa-488 (1:300; ThermoFisher Scientific). The pupal abdominal epidermis was dissected, fixed, and stained with anti-Svb1s (1:300) and anti-rabbit Alexa-488 (1:300; ThermoFisher Scientific) as described in http://gompel.org/wp-content/uploads/2015/09/2003-12-pupal_epidermis.pdf.

### ATAC-seq in pupal abdominal epidermis

For each replicate (n=3) we removed abdominal cuticles from 50 pupae 38-45 hAPF. We carefully removed internal organs with forceps in cold PBS, to retain only epidermal cells, which are attached to the cuticle. We followed the Omni-ATAC protocol (Corces et al. 2017) but using a lysis buffer based on IGEPAL detergent (Lysis Buffer: 10 mM Tris-HCl, pH=7.5, 10 mM NaCl, 3 mM MgCl2, 0.1% IGEPAL CA-630). Library concentration was quantified with a KAPA Library Quantification Kit (Roche) and quality control was performed with the High-Sensitivity DNA Analysis kit in a Bioanalyzer 2100 (Agilent). Libraries were sequenced on a NextSeq 550 system (Illumina) with 37 bp PE reads.

### CUT&RUN against H3K27ac in pupal abdominal epidermis

For each replicate (n=2) we removed abdominal cuticles from 10 pupae 38-45 hAPF. We generated two replicates for the histone H3K27ac antibody (39134, Active Motif) and one control replicate for the Normal Rabbit IgG antibody (Cat. 2729S, Cell Signaling). Both antibodies were used at a 1:100 dilution. After dissection, we washed pupal abdomens by replacing PBS with 1ml of Wash+ buffer (20 mM HEPES pH 7.5, 150 mM NaCl, 0.5 mM Spermidine with Roche complete EDTA-free protease inhibitor) and centrifuging twice at 12000 g for 5 minutes. We resuspended pupal abdomens in 15 ul of BioMag Plus Concanavalin-A-conjugated magnetic beads (ConA beads, Polysciences Inc.) and followed a CUT&RUN protocol for *Drosophila* tissues (https://dx.doi.org/10.17504/protocols.io.umfeu3n). Libraries were prepared using the NEBNext® Ultra™ II DNA Library Prep Kit for Illumina (NEB) following instructions, but with the following changes: (i) adaptors were diluted 1:10 in water for adaptor ligation (step 2), (ii) the size selection of the adaptor-ligated DNA in step 3A was omitted (we proceeded directly to step 3B) and (iii) we performed 14 cycles of PCR with 10 seconds of annealing/extension for enrichment of short DNA fragments. Libraries were sequenced on a NovaSeq 6000 system with 150 bp PE reads.

### ChIP-seq against H3K27ac in svb+ cells of the embryo

Stage 14-15 embryos from a line containing E10::GFP and 7::DsRed transgenes (Preger-Ben Noon et al. 2016) were cross-linked, dissociated and isolated nuclei were immunostained with anti-GFP and anti-DsRed antibodies and the appropriate secondary antibodies. The E10::GFP and 7::DsRed nuclei, which constitute the majority of *svb* expressing nuclei, were then isolated by fluorescence activated cell sorter (FACS). Chromatin from 250,000 nuclei of each cell sub-populations (n=3) was isolated and used for ChIP with anti-H3K27ac and anti-H3 antibodies (Abcam) using the iDeal ChIP-seq kit (Diagenode). Libraries were prepared using the Ovation Ultralow V2 DNA-Seq library preparation kit (NuGen) according to the manufacturer instructions. Libraries were sequenced on a NextSeq 550 system (Illumina) with 50 bp SE reads.

### Data availability

Raw and processed data generated in this paper are available in Gene Expression Omnibus under accession number ###.

### Source of ATAC-seq data from embryonic epidermal cells, wing imaginal disc, eye-antenna imaginal disc, larval brain and T2 pupal leg

For embryonic epidermal cells, the pooled single-cell ATAC-seq fastq files were downloaded from the Gene Expression Omnibus (accession number GSE101581). We filtered reads from epidermal cells from 10-12 hr. embryos (stage 14-15) (Cusanovich et al. 2018)) using files available at https://descartes.brotmanbaty.org/bbi/fly-chromatin-accessibility and https://github.com/shendurelab/fly-atac. Raw fastq files from ATAC-seq experiments from wing imaginal disc, eye-antenna imaginal disc, 3rd instar larva brain and pupa T2 leg were downloaded from Gene Expression Omnibus (accession numbers GSE102841, GSE59078, GSE102441 and GSE113240, respectively).

### Mapping and analysis of NGS data

The fastq files were processed with a custom python pipeline (available at https://github.com/laiker96/fastq_to_bam). Briefly, we ran bbduk.sh (sourceforge.net/projects/bbmap/) to trim adapters and aligned adapter-corrected reads to the *D. melanogaster* reference genome dm6 with BWA (Li and Durbin 2009). Alignment files were filtered based on MAPQ values (>=20) and sam flags with samtools (http://www.htslib.org), and deduplicated with Picard MarkDuplicates from the Picard suite (https://broadinstitute.github.io/picard/). Pearson correlation coefficients between replicates, computing read counts in 1 kb windows, were between 0.94-0.99 for ATAC-seq of pupal epidermis, between 0.79-0.90 for H3K27ac ChIP-seq of *svb* positive cells and 0.96 for H3K27ac CUT&RUN of pupal epidermis. We merged bam files corresponding to the H3K27ac ChIP-seq of E7 cells and E10 cells of the embryo after removing duplicates and subsampling to the size of the smaller dataset. To compare chromatin aperture between contexts, we normalized ATAC-seq experiments taking into account differences in the signal-to-noise ratio between different libraries using S3norm (Xiang et al. 2020). Because we used data from both paired-end and single-end sequencing experiments, we created raw count bedgraph files corresponding to the regulatory region of *svb* using only the first mapped mate from paired-end experiments and used these files as input for the S3norm normalization. The S3 normalized bedgraph files were converted to bigwig files with bedGraphToBigWig (Kent et al. 2010). We fed filtered BAM files to MACS2 (Zhang et al. 2008) to call peaks in ATAC-seq data. For pupal ATAC-seq data, we merged bam files from the different replicates. MACS2 was used with the following parameters: -g dm -f BAM --keep-dup all --shift -50 --extsize 100 --nomodel. To normalize H3K27ac data we used IgG (CUT&RUN) and histone H3 (ChIP-seq). H3K27ac experiments were normalized with bamCompare from the DeepTools suite (Ramírez et al. 2016) with --scaleFactorsMethod SES and a bin size of 1. Genome browser plots were generated with the pyGenomeTracks package (Lopez-Delisle et al. 2021).

### X-GAL stainings

Third-instar larvae were dissected in PBS and fixed in PBS with 4% formaldehyde for 10 minutes. Staged pupae were removed from the pupal case and then fixed in PBS with 4% formaldehyde for 15 minutes. After washing in PBT (PBS + 0.1% Triton X-100), samples were incubated with X-Gal solution (5 mM K4[Fe+2(CN)6], 5 mM K3[Fe+2(CN)6], 1 mg/ml X-Gal in PBT) at 37 C for 1 hr. The samples were mounted and imaged with bright-field microscopy. We used a fly line that does not carry a *lacZ* reporter as negative control.

### Microscopy and Image Analysis

Pupae of the desired stages were removed from the pupal case and placed in a microscope slide for imaging. For live GFP imaging in pupa, GFP signal was measured over a z stack in a confocal microscope. Images were analyzed using ImageJ software (http://rsb.info.nih.gov/ij/). We used the sum projection of the z stacks to analyze qualitatively the GFP levels between different stages. To analyze the effect of enhancer deletions in *svb*BACs we measured GFP and DsRed levels in pupae carrying *svb*BAC-GFP (wild-type and deletions) and svbBAC-DsRed (wild-type). GFP and DsRed signals were measured sequentially over a z stack in a confocal microscope. Images were analyzed using ImageJ software (http://rsb.info.nih.gov/ij/). First, background was subtracted using a 50-pixel rolling-ball radius in each slice of the confocal z stack. Then, we calculated the sum projection of the z stacks for each channel to compare GFP versus DsRed levels. Segmentation masks were applied with Ilastik 1.2.0 software (http://ilastik.org) to the sum projections of the GFP channel. We measured the fluorescence mean intensities of each nucleus with the ‘‘Analyze particles’’ tool in ImageJ. Then, we calculated the average of the fluorescence mean intensity of all segmented nuclei. Last, we computed the ratio GFP/DsRed in each nucleus and calculated the average ratio for all segmented nuclei.

Supplementary Figure 1. Open chromatin profile of the *svb* regulatory region in different developmental contexts. The position of the embryonic enhancers of *svb* is shown below.

Supplementary Figure 2. X-Gal stainings of whole pupae for each reporter line.

Supplementary Figure 3. Detection of Svb in imaginal discs, brain and foregut of the larva through immunofluorescence. Colocalization between GFP (*svb*BAC-GFP expression) and Svb (red).

Supplementary Figure 4. X-Gal stainings of larval tissues (epidermis, foregut and brain) for each reporter line. N.E.: No expression.

Supplementary Figure 5. Odds ratios of conserved and non-conserved bases in regions of open chromatin for the chosen regulatory regions (A) and the 10,000 random windows of the *Drosophila* genome (B).

Supplementary Table 1. Coordinates and fraction of bases in conserved elements for the different DNA regions within *svb*.

Supplementary Table 2. List of primers used for recombineering.

Supplementary Table 3. List and sources of the ATAC-seq data used for the analysis of Figure 7.

