## Supplementary figures and images for "The density of regulatory information is a major determinant of evolutionary constraint on non-coding DNA in *Drosophila*"

### Supplementary Figures

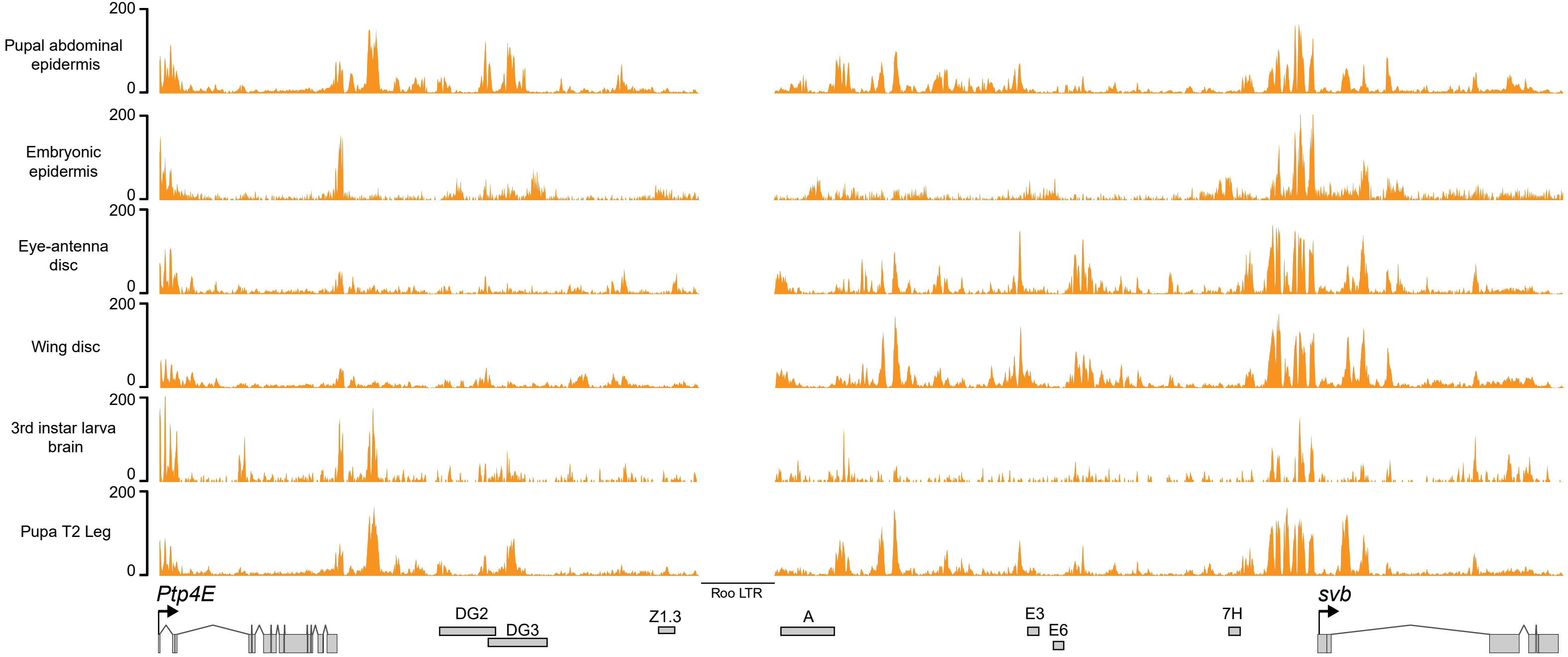

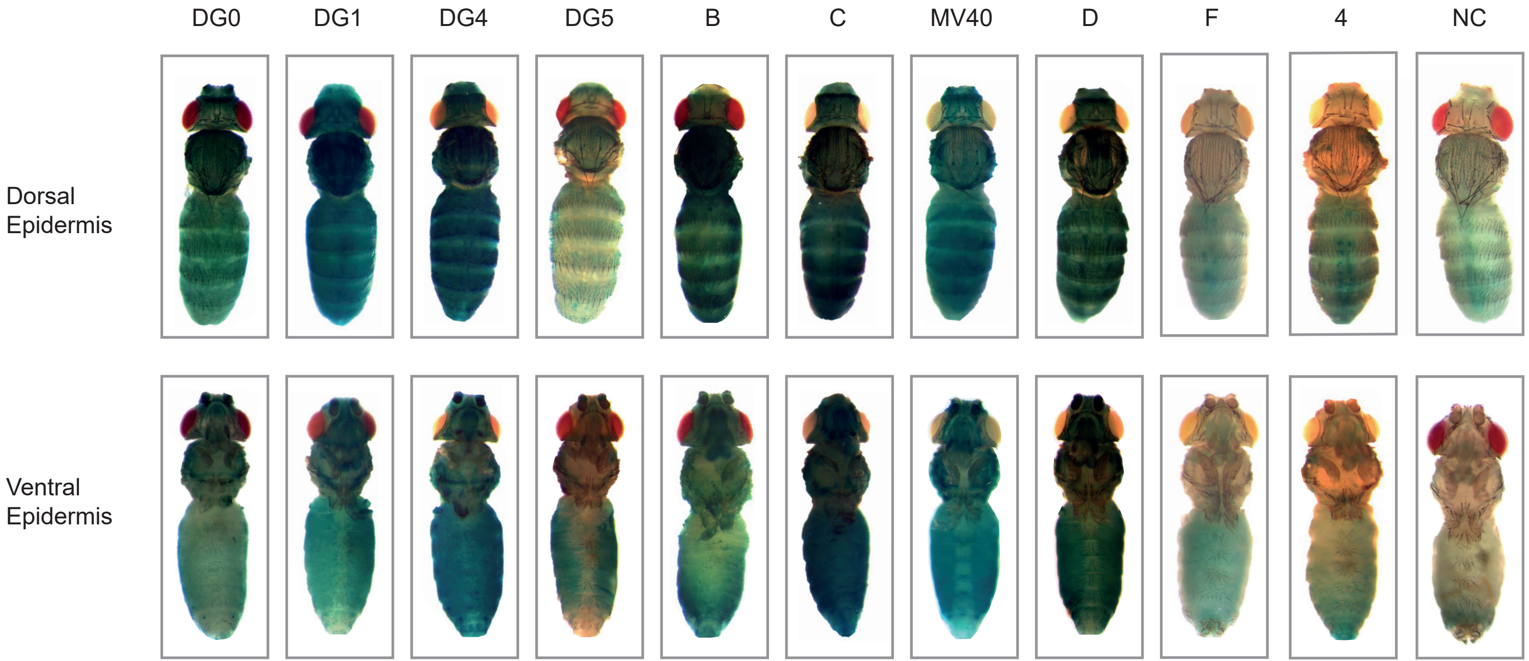

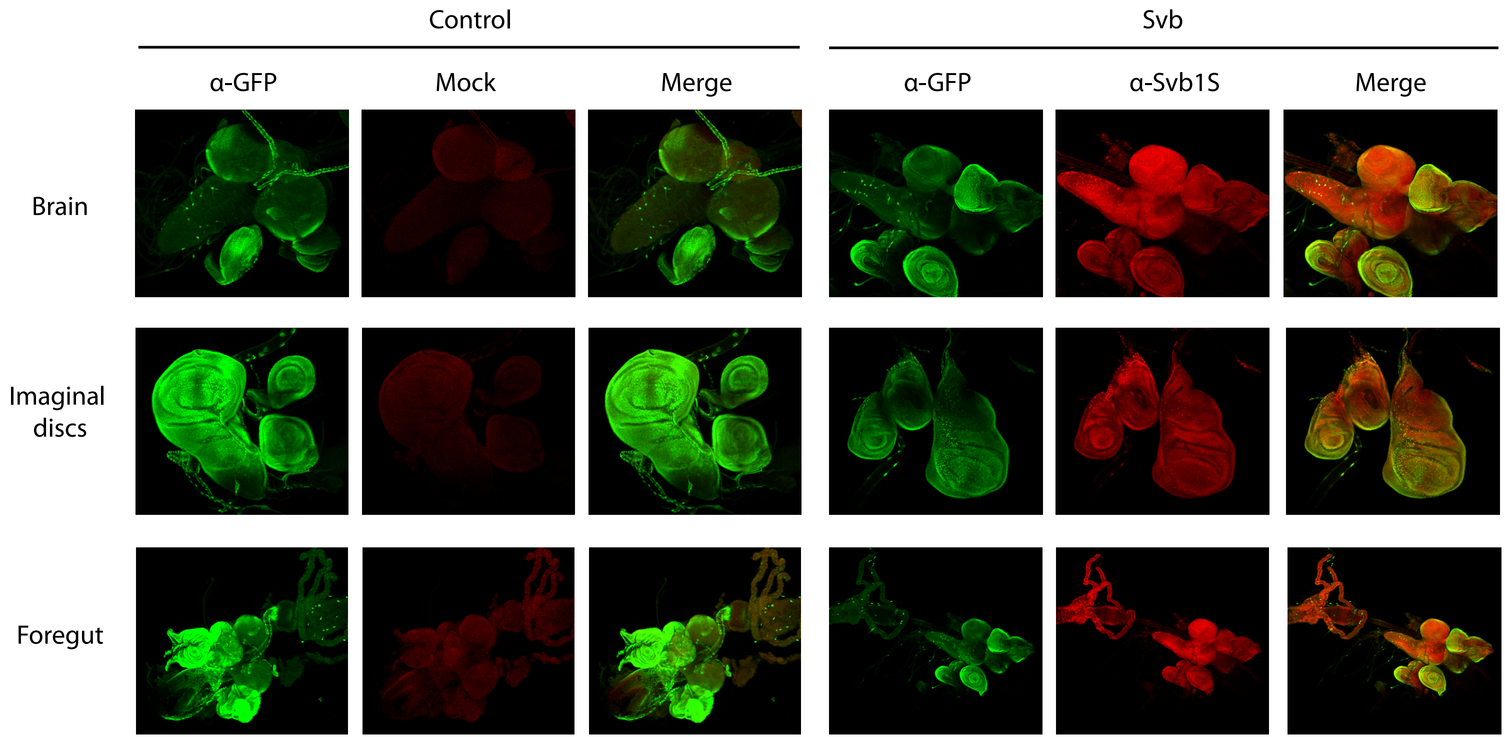

Figure S4

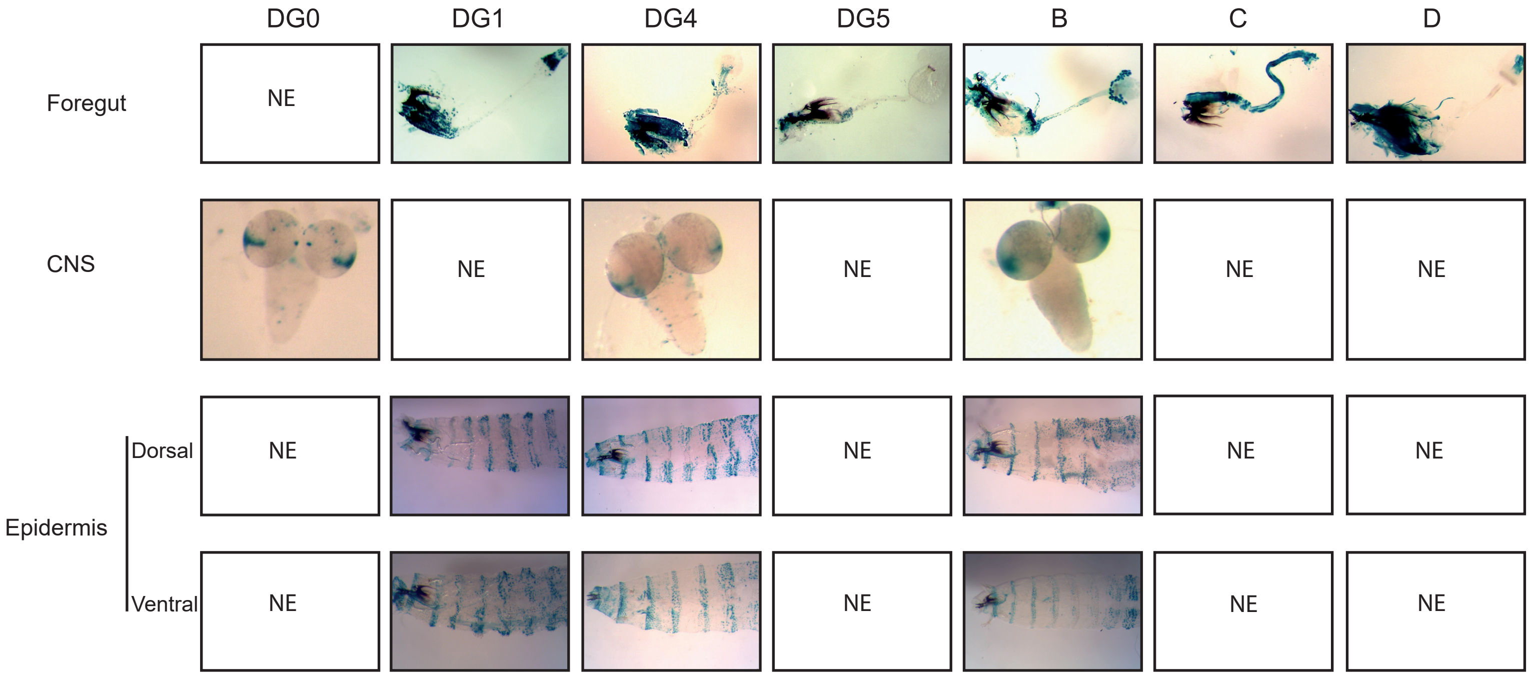

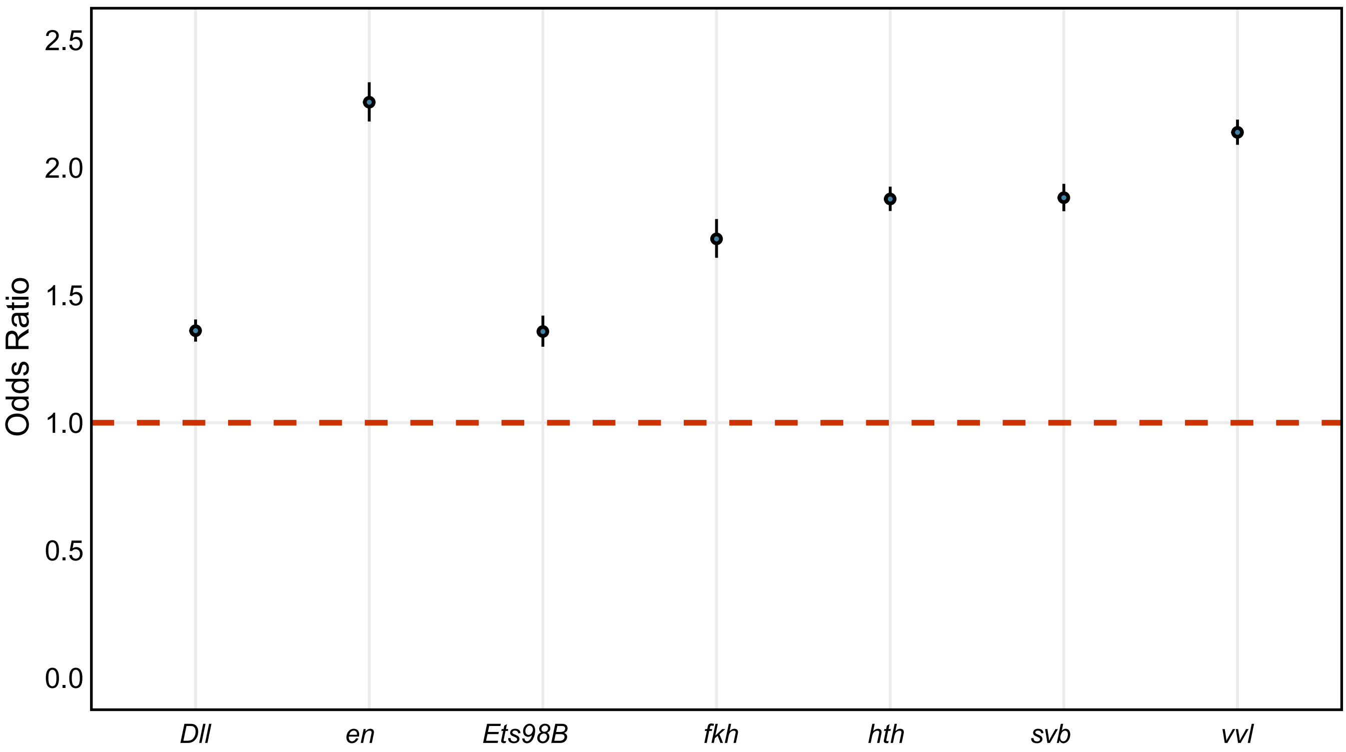

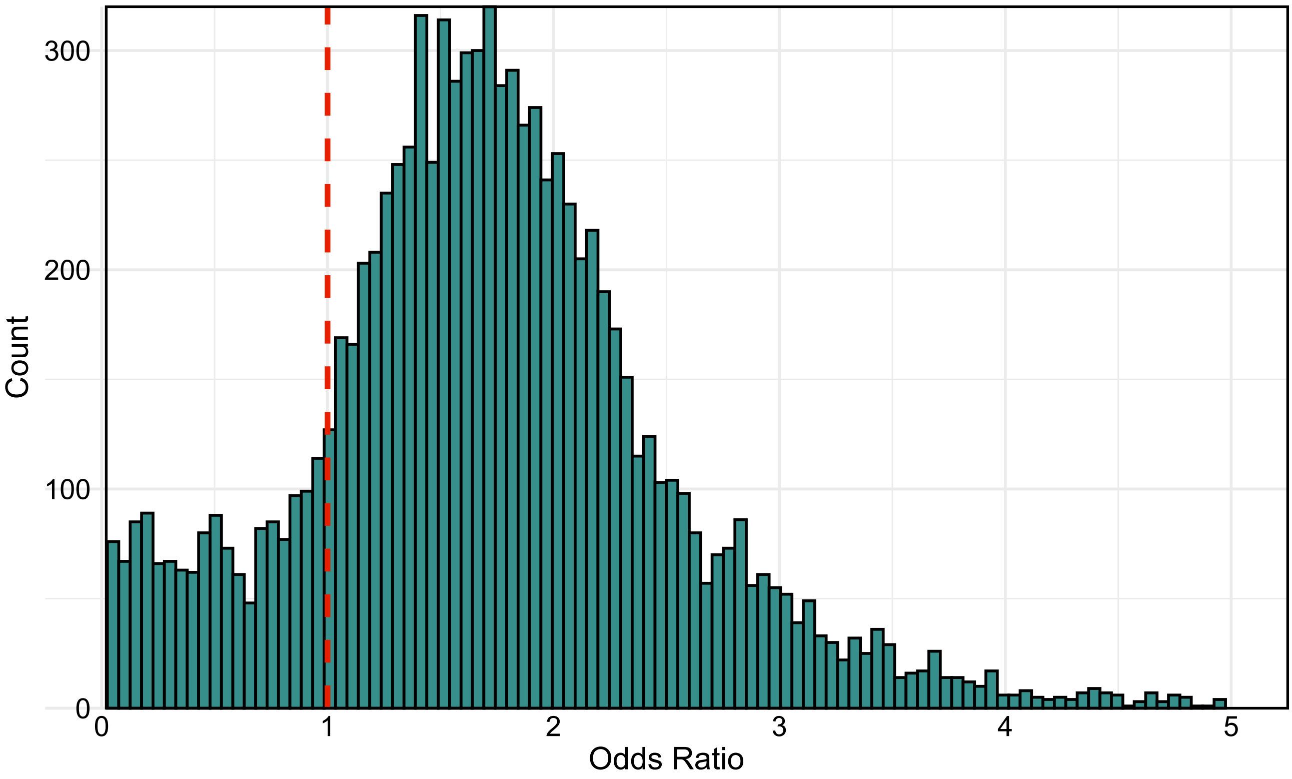
